# Farming practices in the landscape have an impact on grassland arthropods, in addition to local conditions and land cover

**DOI:** 10.1101/2024.11.25.625040

**Authors:** Théo Brusse, Jodie Thénard, Ronan Marrec, Gaël Caro

## Abstract

Arthropods deliver vital ecosystem services in agricultural landscapes, but their diversity is threatened by intensified farming and landscape simplification. Despite this, the relative impacts of farming practices at the landscape level remain underexplored compared to local and land cover effects. In this study, we dissect the influence of local and landscape factors, including farming practices, on arthropod communities. By sampling 18 grasslands in northeastern France during spring and autumn 2021, we collected 14 arthropod families, identifying all species of carabids and spiders. We conducted surveys of farming practices to estimate their intensity at landscape level. Our results highlight that environmental factors affect arthropod communities differently between seasons. Intensity of farming practices at landscape level was less explanatory of arthropod communities, but this impact is additional to that of local conditions and land cover. Our study underscores the critical need to integrate farming practices into landscape-level management strategies to safeguard arthropod diversity.

## 1. Introduction

Increased demand for food has led to changes in farming practices and landscapes, such as the reduction and fragmentation of semi-natural habitats, the increase in the size of cultivated plots and farms, and the increased use of fertilizers and pesticides (Benton et al., 2003; Robinson and Sutherland, 2002; Tilman et al., 2002). One of the consequences is the decline in biodiversity over the last 70 years, which biodiversity is essential to the functioning and resilience of ecosystems (Meier et al., 2022; Robinson and Sutherland, 2002; Tilman et al., 2002). One way of preserving biodiversity in intensive agricultural landscapes is to maintain areas of semi-natural habitats (Garibaldi et al., 2021; Tscharntke et al., 2021), including grasslands, which are recognized as a particularly important habitat for biodiversity sustainability by providing feeding and nesting resources (Petermann and Buzhdygan, 2021). Among threatened taxa, grassland arthropods are of particular interest as they play an important role in providing and maintaining ecosystem functions, both for the grassland and for the surrounding (agro-)ecosystems (Bardgett and Van Der Putten, 2014; Losey and Vaughan, 2006).

Arthropod diversities are impacted directly and indirectly by local management practices (Rusch et al., 2010). These practices, such as mowing, grazing, and fertilization, drive directly local plant community diversity and functional structure (Prather et al., 2020; Socher et al., 2013; Van Klink et al., 2015), and consequently, arthropod communities inhabiting these grasslands. In fact, plant development affects the biomass of arthropods, which is higher in developed and diverse grasslands, characterized by a high Normalized Difference Vegetation Index (NDVI), because of the increased availability of resources and the resulting diversification of food webs (Ebeling et al., 2018; Fernández-Tizón et al., 2020). In addition, NDVI value is related to near-ground temperature in grassland (Maclean et al., 2019) which influences arthropod activity. Thus, warmer grasslands in winter favor their presence by providing favorable shelter for arthropods during overwintering (Seibold et al., 2016) and consequently encourage their activities throughout the season.

One of the recognized advantages of grasslands in agricultural landscapes is that they can serve as reservoirs and shelters for many species from other patches. The degree of grassland accessibility determines the influence of surrounding landscape on local arthropod abundance and diversity through dilution/concentration processes (Vasseur et al., 2013). The known landscape drivers considered to be the main ones explaining arthropod diversity are linked to the heterogeneity of land cover, whether in terms of composition (e.g., quantity of semi-natural habitats, crop diversity) or configuration (e.g., mean field size) (Estrada-Carmona et al., 2022; Marja et al., 2022; Tscharntke et al., 2021).

Nevertheless, characterizing landscape heterogeneity only through land cover is limited, and does not integrate the management intensity of landscape elements, despite its widely recognized impact at field level (Rusch et al., 2010). However, the number of studies on the effect of farming practices at landscape level is very limited, although some have shown landscape-wide intensity influences the abundance-activity of arthropod communities (Brusse et al., 2024a). Even fewer studies have considered the configuration of farming practices in the landscape. Apart from one study on the impact of tillage configuration on surface runoff in vineyards (Colin et al., 2012), none, to our knowledge, has used the configuration of practices to explain biodiversity patterns. Yet, the configuration and the degree of intensity of farming practices can in fact modify the relative quality of fields of the same or different crops, and thus alter their attractiveness to organisms or their ability to promote/limit their movement in the landscape (Herzog et al., 2006; Marrec et al., 2022; Maudet et al., 2024).

To make agricultural landscapes favorable to biodiversity — and therefore to arthropods —, it is essential to consider both the traditionally used parameters of land cover and local conditions (Estrada-Carmona et al., 2022; Tscharntke et al., 2021), and parameters of intensity and configuration of farming practices at landscape level, and quantify their relative effects. This will make it possible to determine whether farming practices at landscape level have a greater or lesser influence than parameters recognized as important for arthropods (Estrada-Carmona et al., 2022; Tscharntke et al., 2021), and whether it is relevant to include them in future studies. Our study therefore focused on a network of 18 permanent grasslands for which we characterized the local parameters and the landscape parameters belonging to both the heterogeneity of land cover and farming practices. The aim of our study was to study the effect of local and landscape parameters on arthropod communities, by considering the intensity and configuration of farming practices in the landscape. Then, to determine the importance of farming practices at landscape level in explaining patterns of arthropod diversity, we aimed to estimate the proportion of variance that they explain compared to the variance explained by the other local and landscape parameters traditionally considered in studies.

## 2. Material and methods

### 2.1. Study area

The study was carried out in the Parc Naturel Régional de Lorraine in northeastern France, covering an area of 680 km2 (48°48′N, 6°43′E) (Fig. 1a). This area was selected because grasslands are embedded in a mosaic of crops, forests, and urban areas, and are subject to various intensities of farming practices at the landscape level. The region experiences a temperate climate, with an average yearly rainfall of 765.2 mm and an annual mean temperature of 10.6 °C. Eighteen permanent grasslands were chosen within the study area to capture a range of landscape openness levels (i.e., based on the percentage of woody elements within a 500-meter radius buffer around the grassland center) (see Fig. 1b). Given that the area is primarily used for grassland and crop production, these account for 45.2% (standard deviation = 20.5%) and 29.7% (sd = 23.1%) of the landscapes selected, respectively. Forests make up 18.7% (sd = 29.9%), water bodies 4.5% (sd = 10.7%), and urban areas 1.9% (sd = 6.1%) of these landscapes.

**Figure 1.**
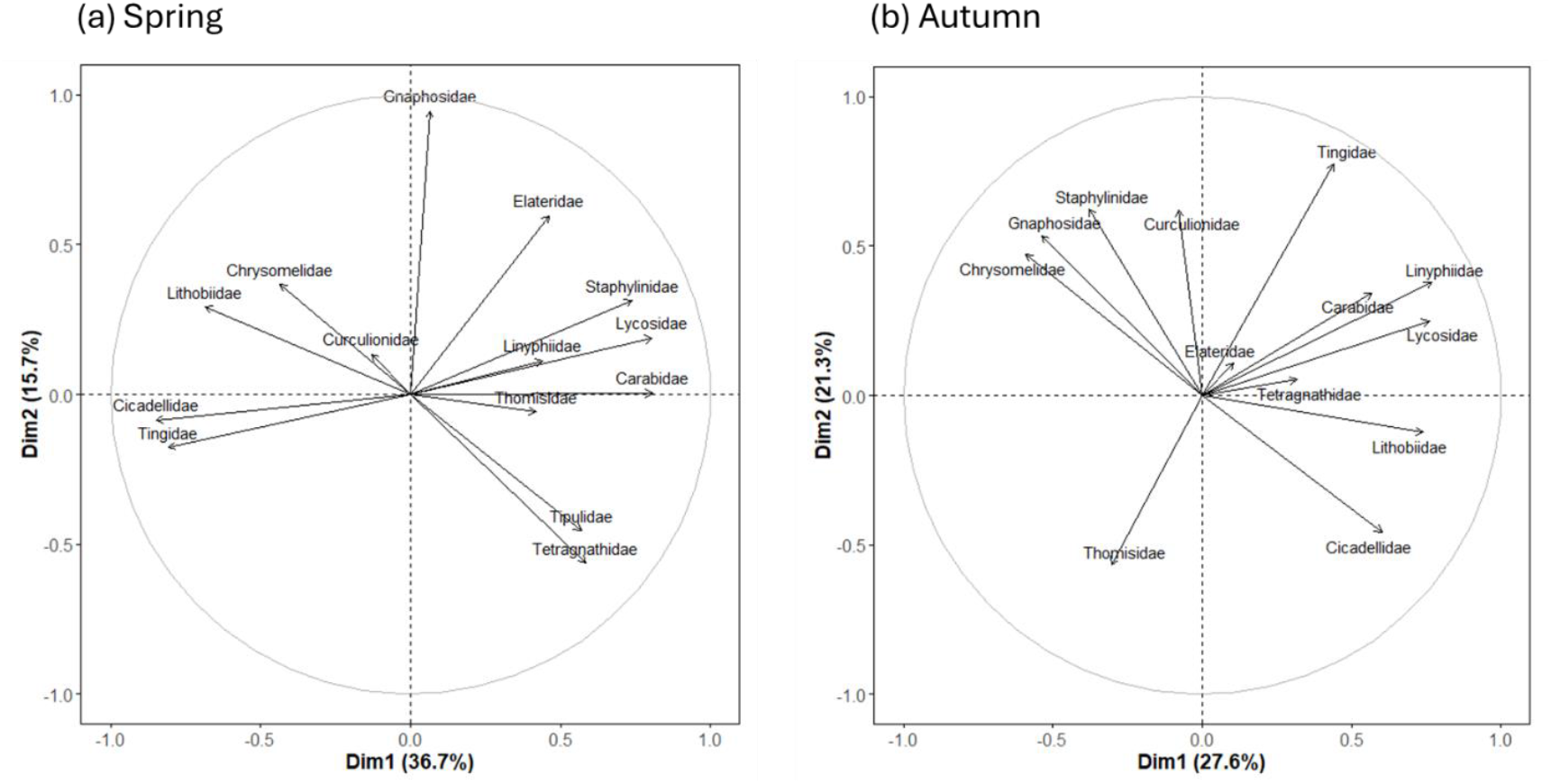
Results of principal component analysis (PCA) illustrating composition of arthropod communities (a) in spring and (b) in autumn.

### 2.2. Arthropod sampling

Arthropods were collected twice annually in 2021 from 18 chosen grasslands using pitfall traps, with sampling occurring once in spring (May) and again in autumn (September). These traps were employed specifically to capture ground-dwelling arthropods (Luff, 1975). Five pitfall traps (⌀ = 10 cm; height = 15 cm) were systematically installed at the center of each grassland, spaced 15 meters apart, to sample organisms adapted to local habitat conditions representatively. Each trap was filled to one-third of its capacity with a white vinegar solution and protected from rain by a cover, and traps were left undisturbed in the field for a period of seven days.

A taxonomic specialist identified the ground-dwelling arthropods at the family level. For analysis, we focused on families that represented at least 95% of overall activity-abundance: Gnaphosidae, Linyphiidae, Lycosidae, Tetragnathidae, Thomisidae, Staphylinidae, Carabidae, Lithobiidae, Chrysomelidae, Curculionidae, Elateridae, Cicadeliidae, Tingidae, and Tipulidae. Family richness was used to assess arthropod community diversity. In addition, species-level identification was performed for two taxonomic groups: carabid beetles (Coulon et al., 2011; Jeannel, 1942, 1941) and spiders (Nentwig et al., 2024). The TaxRef (2024) framework served as the taxonomic reference for invertebrates. For each grassland community and for each season, we calculated (1) the absolute activity-abundance of all arthropods, carabids, and spiders, (2) family richness for arthropods and species richness for carabids and spiders (i.e., the number of families or species per grassland), (3) family and species evenness based on Pielou’s index, and (4) the community composition of all arthropods, carabids, and spiders through Principal Component Analysis (PCA) using the FactoMineR R package (Lê et al., 2008). For PCA, the activity-abundance of each family or species was used. The optimal number of PCA dimensions was determined using the nFactors R package (Raiche and Magis, 2022), with each dimension reflecting different levels of correlation with family or species abundance-activity.

### 2.3. Environmental data

#### 2.3.1. Local parameters

The grasslands sampled were characterized by three parameters: the Normalized Difference Vegetation Index (*NDVI*), the *temperature near the ground*, and the *intensity of farming practices*. First, we calculated NDVI as a proxy of vegetation vigor and quantity (Table 1). NDVI is a key indicator of arthropod biomass in grassland habitats (Fernández-Tizón et al., 2020). We calculated the NDVI for April and September 2021 (dates corresponding to arthropod sampling periods) using multi-spectral images from the Sentinel-2 satellite, utilizing the red (R) and near-infrared (NIR) bands with the formula: (NIR – R) / (NIR + R). Second, since previous studies have indicated that arthropod abundance-activity in grasslands is influenced by temperatures near the ground (Seibold et al., 2019), we estimated the average monthly temperature under vegetation near the soil surface for April and September 2021 using the *microclima* R package (Maclean et al., 2019). This mechanistic modeling approach employs a wide range of parameters (including NDVI, vegetation cover, macroclimatic temperature, and atmospheric conditions) to generate maps of predicted microclimatic temperature at the scale of the sampled grasslands (see Table S1 for a complete list of parameters and their sources). Finally, for farming practices implemented at the sampled grassland level, we surveyed farmers about the number of mowings, mowing height, livestock load, and the quantity of nitrogen input. From these parameters, we calculated the overall intensity value of the sampled grasslands, according to the formula proposed by Herzog *et al*. (2006).

**Table 1.** List of environmental parameters used for explained patterns of arthropod communities.

| Category | Variable | Extent |
| --- | --- | --- |
| Sampled grassland parameters (local) | NDVI | 0.59 - 0.82 |
|  | Temperature near the ground | 10 - 17 °C |
|  | Intensity of farming practices | 0.00 - 0.86 |
| Land cover heterogeneity (landscape) | Proportion of grassland in the landscape | 6-78 % |
|  | Crop diversity | 1-11 |
|  | Mean field size | 3 - 13 ha |
| Resources and<br>microclimate<br>(landscape) | NDVI-landscape | 0.40 - 0.76 |
|  | Mean wood grain | 0.03 - 0.97 |
| Farming<br>practices<br>(landscape) | LWI (landscape-wide<br>intensity) | 0.13 - 0.44 |
|  | a-LWI (aggregation of<br>LWI) | 88 - 98 |
|  | d-low-I (mean distance<br>between low-intensity<br>patches) | 0 - 600 m |

#### 2.3.2. Landscape parameters: land cover heterogeneity

Three parameters were used to describe land cover heterogeneity at the scale of 500-m radius landscape buffer centered on the centroid of sampled grasslands: *proportion of grassland in the landscape, crop diversity*, and *mean field size* (Table 1). We chose these three parameters because they are acknowledged as significant factors for agricultural biodiversity (Estrada-Carmona et al., 2022; Marja et al., 2022; Tscharntke et al., 2021) and encompass most aspects of land cover heterogeneity (Fahrig et al., 2011). The French graphic parcel register (https://geoservices.ign.fr/rpg) was used to record the identity of crops in the landscape and then derive the proportion of grasslands in the landscape, land cover diversity and mean field size.

#### 2.3.3. Landscape parameters: resources and microclimate in the landscape

The mean value of NDVI within 500-m radius landscape buffer (*NDVI-landscape*) was used as a proxy of resource continuity at landscape level (Iuliano and Gratton, 2020) (Table 1). As with the method for calculating NDVI at the scale of the sampled grassland, we calculated the *NDVI-landscape* for April and September 2021 using multi-spectral images from the Sentinel-2 satellite. Then, we calculated the *mean wood grain* within landscape buffer as a measure of landscape openness and as a proxy of microclimatic effects of surrounding landscape elements (e.g., wind-break effect, humidity retention) (Tougeron et al., 2022) (Table 1). The value of the wood grain was calculated for each 50 × 50 m pixel within the 500 m concentric landscape buffers, and accounts for both the composition and the configuration of woody elements. The value of each pixel in the 500 m buffer was calculated from the woodland structure within 400 m of that pixel. The value of each pixel ranges from 0 to 1, depending on the density of woodland within a 400 m radius of the pixel. Low values of wood grain correspond to a high density of woody elements, and high values in landscapes to low densities of woody elements (Vannier et al., 2011). Then, the *mean wood grain* was calculated per landscape by averaging the value of wood grain in each pixel inside the 500 m concentric landscape buffers.

#### 2.3.4. Landscape parameters: farming practices

We calculated three parameters representing the heterogeneity of farming practices within 500-m radius landscape buffer: landscape-wide intensity (*LWI*), aggregation of landscape wide intensity (*a-LWI*), and mean distance between low-intensity patches (*d-low-I*) (Table 1).

We followed the method developed by Maudet et al. (2024) to estimate *LWI*. This PCA-based method allows to circumvent the need for a subjective and limited selection of practices and integrate potentially correlated variables and reduce the number of dimensions to be considered. Agricultural practices applied in 2021 within grasslands and crop fields located within a 500-meter radius from the core of the sampled grasslands were documented through standardized farmer surveys (Table S2). A total of 39 farmers from 18 different landscapes participated, providing data on 180 fields. The survey covered, on average, 91.4% (SD = 9.9%) of the agricultural land in each landscape. To minimize the cumulative impact of multiple farming practice variables while preserving as much information as possible from the original data, we summarized each recorded variable through a principal component analysis (PCA) (see Figure S1). An intensity index, following Herzog et al. (2006), was then calculated, which uses the selected principal components (PCs) as combined indicators and applies their eigenvalues as weights, following this equation:

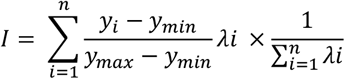

where *I* was the intensity index, *y*_*i*_ the observed value for the *PC*_*i*_ and *λ*_*i*_ its eigenvalue, *y*_*min*_ and *y*_*max*_ the minimum and maximum observed value on *PC*_*i*_. The number of PCs to retain was estimated according to Cattell-Nelson-Gorsuch procedure, using the *nFactors* R package (Raiche and Magis, 2022). Finally, we averaged the intensity values of each field in the landscape, weighted by their surface area, to obtain the value of *LWI*.

In addition to *LWI*, we also considered the configuration of intensity in the landscape (*a-LWI* and *d-low-I*). As the intensity values for each field are quantitative, it is necessary to convert them into qualitative classes to calculate the two configuration parameters. We carried out a sensitivity analysis revealing that dividing the intensity into eight equal intensity classes maximized the diversity of intensity patches while moving away from the land cover categorization (Figure S2). The first configuration index (*a-LWI*) indicates the level of grouping between the different intensity classes. *Aggregation* equals 0 for maximally disaggregated and 100 for maximally aggregated intensity classes (Hesselbarth et al., 2019). The second index (*d-low-I*) corresponds to the average Euclidean distance between patches of lower intensity. The index approaches 0 as the distance to the nearest neighbor decreases (i.e., patches of low intensity class are more aggregated), and increases, without limit, as the distance between neighboring patches of low intensity class increases (i.e., patches are more isolated) (Hesselbarth et al., 2019).

### 2.4. Statistical analyses

All analyses were performed using R v.4.1.2 (R Core Team). Because of the large number of independent parameters that could influence arthropods, all were not included in a single model. Rather, we utilized a hierarchical approach to test the response of arthropod community response to environmental factors (Caro et al., 2016). We modelled separately, and for each taxonomic group (carabids, spiders, and all arthropods), abundance-activity (log-transformed), species/family richness (log-transformed), and community composition as the response variables in linear models (LM). We constructed models following four steps, each step complexifying the model and testing the effect of a set of specific parameters. At each step, the best model was selected by comparing AIC values (ΔAIC < 2) between all possible sub-models using a stepwise deletion procedure and the combination of parameters of this best model was retained for the next step. For each selected model, we verified the level of collinearity in our model by calculating the variance inflation factor (VIF) for each variable (Smith et al., 2009). Collinearity refers to the degree of correlation between two or more explanatory parameters in a regression model and can affect the accuracy and stability of the estimates. We used the threshold of VIF < 3 to indicate low collinearity (Smith et al., 2009). The procedure of model construction is described below.

As a first step, *NDVI, temperature near the ground*, and *intensity of farming practices* were included to the model to test effects of sampled grasslands factors. In step 2, land cover parameters (i.e., *proportion of grassland in the landscape, crop diversity*, and *mean field size*) were added to the resulting model of step 1. In step 3, microclimatic and resource continuity in the landscape (i.e., *mean wood grain* and *NDVI-landscape*) were added to the parameters selected in step 2. Finally, in step 4, *LWI, a-LWI* and *d-low-I* were added to the resulting model of step 3 to characterize the intensity and configuration of farming practices at the landscape level.

Then, we distinguished the portion of variance explained by farming practices in the landscape from the variance explained by the local parameters and the land-cover parameters. For each biological variable to be explained, we extracted (i) the R^2^ value of a model including all the previously selected farming practice parameters at the landscape level (parameters selected during step 4), (ii) the R^2^ value of a model including local parameters (parameters selected during step 1), (iii) the R^2^ value of a model including parameters selected during steps 2 and 3, and (iv) the R^2^ values of each combination of models possible from these first three models. Based on these R^2^ values, we used the *varPart()* function in the ModEva R package (Barbosa *et al*. 2016)to estimate the proportion of variance explained by farming practices at landscape level, local parameters, and land-cover and resources parameters.

## 3. Results

### 3.1. Description of arthropod communities

A total of 6,558 arthropods were sampled over the two sampling periods, including a total of 1,078 spiders and 1,260 carabids. Arthropod communities were composed of an average of 10.60 taxonomic families (sd = 1.49), 7.66 spider species (sd = 3.98), and 6.19 carabid species (sd = 2.51). Pielou’s equitability index (proxy of evenness) was on average 0.77 (sd = 0.09) for arthropod families, 0.71 (sd = 0.22) for carabid species, and 0.82 (sd = 0.15) for spider species.

We kept the first three dimensions of the PCA to characterize spring arthropod community composition, accounting for 62.2 % of the total variance (Figure 1a). The first dimension explained 36.7 % of the total variance. It contrasted grasslands with communities rich in Carabidae, Lycosidae, and Staphylinidae in its positive part, with grasslands with communities rich in Tingidae, Cicadellidae, and Lithobiidae in its negative part. The second dimension accounted for 15.7 % of the total variance. It contrasted grasslands with communities rich in Gnaphosidae, in its positive part to grasslands with communities rich in Tetragnathidae in its negative part. The third dimension accounted for 11.8 % of the total variance. It was positively correlated with the quantity of Curculionidae in communities. To characterize autumn arthropod community composition, we kept the first three dimensions of the PCA accounting for 62.9 % of the total variance (Figure 1b). The first dimension explained 27.6 % of the total variance. It contrasted grasslands with communities rich in Linyphiidae, Lycosidae, and Lithobiidae in its positive part to grasslands with communities rich in Chrysomelidae in its negative part. The second dimension accounted for 31.3 % of the total variance. It contrasted grasslands with communities rich in Tingidae and Staphylinidae in its positive part to grasslands with communities rich in Thomisidae in its negative part. The third dimension accounted for 14.1 % of the total variance. As for spring, it was positively correlated with the quantity of Curculionidae in communities.

For carabids, we kept the first three dimensions of the PCA to characterize spring community composition, accounting for 46.1 % of the total variance (Figure S3). The first dimension explained 21.0 % of the total variance. It was positively correlated to abundance-activity of *Pterostichus anthracinus* and *Poecilus cupreus*. The second dimension accounted for 13.1 % of the total variance. It contrasted grasslands with communities rich in *Bembidion obtusum* in its positive part, to grasslands with communities rich in *Amara communis* in its negative part. The third dimension accounted for 12.0 % of the total variance. It was positively correlated to abundance-activity of *Carabus auratus*. To characterize carabid community composition in autumn, we kept the first three dimensions of the PCA accounting for 52.5 % of the total variance (Figure S3). The first dimension explained 23.1 % of the total variance. It contrasted grasslands with communities rich in *Pterostichus anthracinus* in its positive part, to grasslands with communities rich in *Harpalus dimidiatus* in its negative part. The second dimension accounted for 16.7 % of the total variance. It contrasted grasslands with communities rich in *Poecilus cupreus* in its positive part to grasslands with communities rich in *Agonum lugens* in its negative part. The third dimension accounted for 12.7 % of the total variance. It contrasted grasslands with communities rich in *Anisodactylus binotatus* in its positive part to grasslands with communities rich in *Pterostichus melanarius* in its negative part.

For spiders, we kept the first three dimensions of the PCA to characterize spring community composition, accounting for 46.5 % of the total variance (Figure S4). The first dimension explained 21.7 % of the total variance. It contrasted grasslands with communities rich in *Pardosa prativaga*, in its positive part to grasslands with communities rich in *Haplodrassus signifier* in its negative part. The second dimension accounted for 13.7 % of the total variance. It contrasted grasslands with communities rich in *Hahnia nava* in its positive part to grasslands with communities rich in *Xysticus kochi* in its negative part. The third dimension accounted for 11.2 % of the total variance. It was positively correlated to abundance-activity of *Pelecopsis parallela*. For characterize spider community composition in autumn, we kept the first three dimensions of the PCA accounting for 64.9 % of the total variance (Figure S4). The first dimension accounted for 34.3 % of the total variance. It was positively correlated to abundance-activity of *Pardosa tenuipes*. The second dimension accounted for 17.4 % of the total variance. It contrasted grasslands with communities rich in *Erigone dentipalpis* in its positive part to grasslands with communities rich in *Troxochrus scabriculus* in its negative part. The third dimension accounted for 13.2 % of the total variance. It was negatively correlated to abundance-activity of *Agyneta rurestris*.

### 3.2. Effects of environmental parameters on arthropod communities

The local parameters were more often selected than landscape ones in the models considering spring communities, indicating a higher relevance of local rather than landscape parameters. (Figure 2a). Among the 18 models, local parameters were selected in 33 % of cases, land-cover parameters in 22 %, and farming practices at landscape level in 17 %. Of all local parameters, the *NDVI* of sampled grasslands is the one that most often influenced biological parameters (significant effect in 33 % of models). Among landscape parameters, although rarely selected in models, *NDVI-landscape* and *mean field size* were those that most often influenced biological variables (significant effect in 22 % and 17 % of models, respectively). Farming practices at the landscape level explained a rather low variance in biological variables compared with other environmental parameters (6 % - 7 %) (Figure 3a). Local parameters explained between 19 and 35 % in average of the variance in biological variables, and land cover parameters between 16 and 22 %.

**Figure 2.**
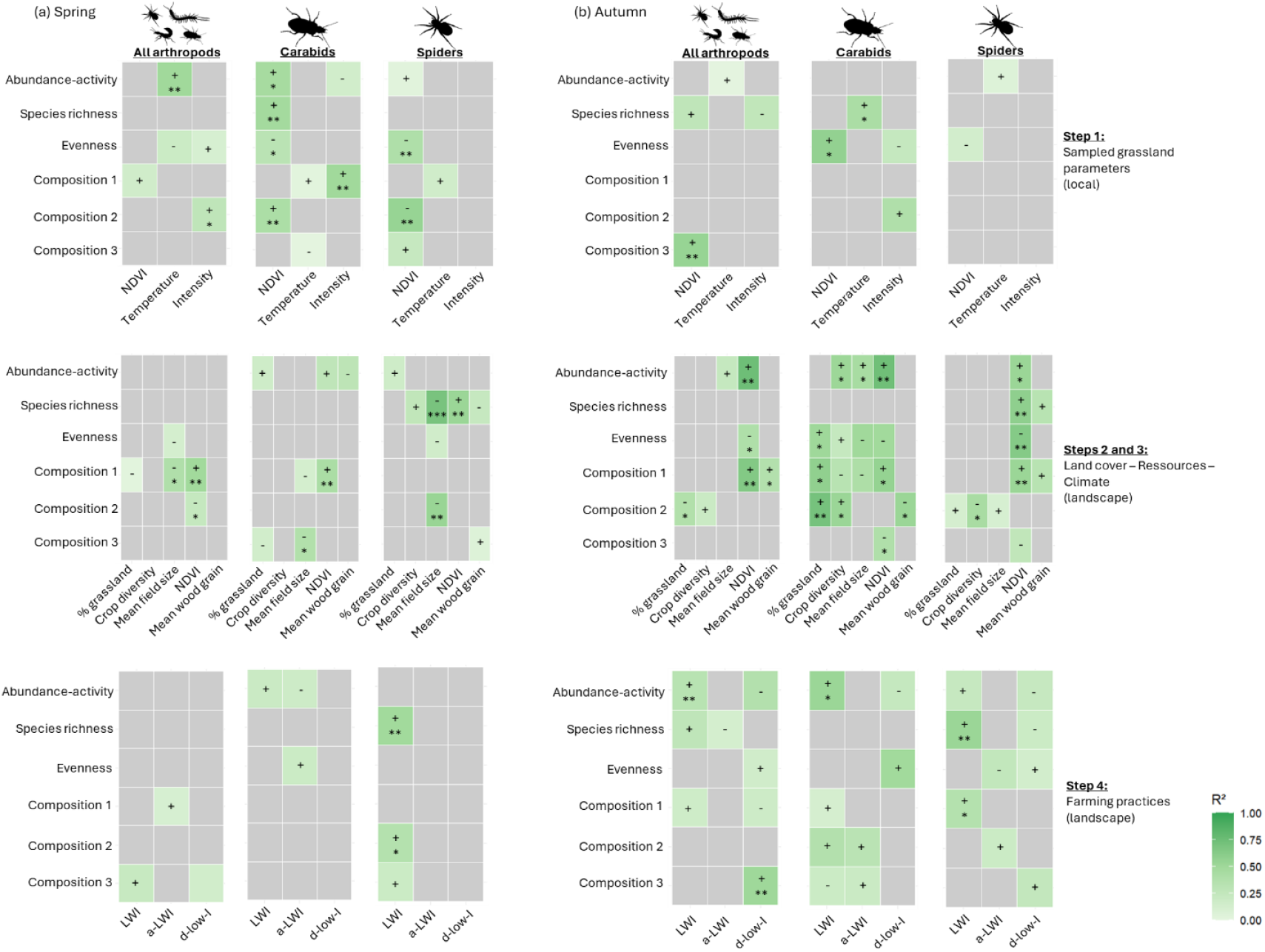
Summary of model selections and effects of selected variables on arthropod communities (a) in spring and (b) in autumn. Grey boxes correspond to parameters not selected during model selection. The degree of green is proportional to the value of the R^2^ of the corresponding variable on the biological variable. The + sign indicates a positive effect of the selected variable on the biological variable, and the - sign a negative effect. Symbols: 0.01 < p-value <0.05 (*); 0.001 < p-value <0.01 (**); p-value <0.001 (***).

**Figure 3.**
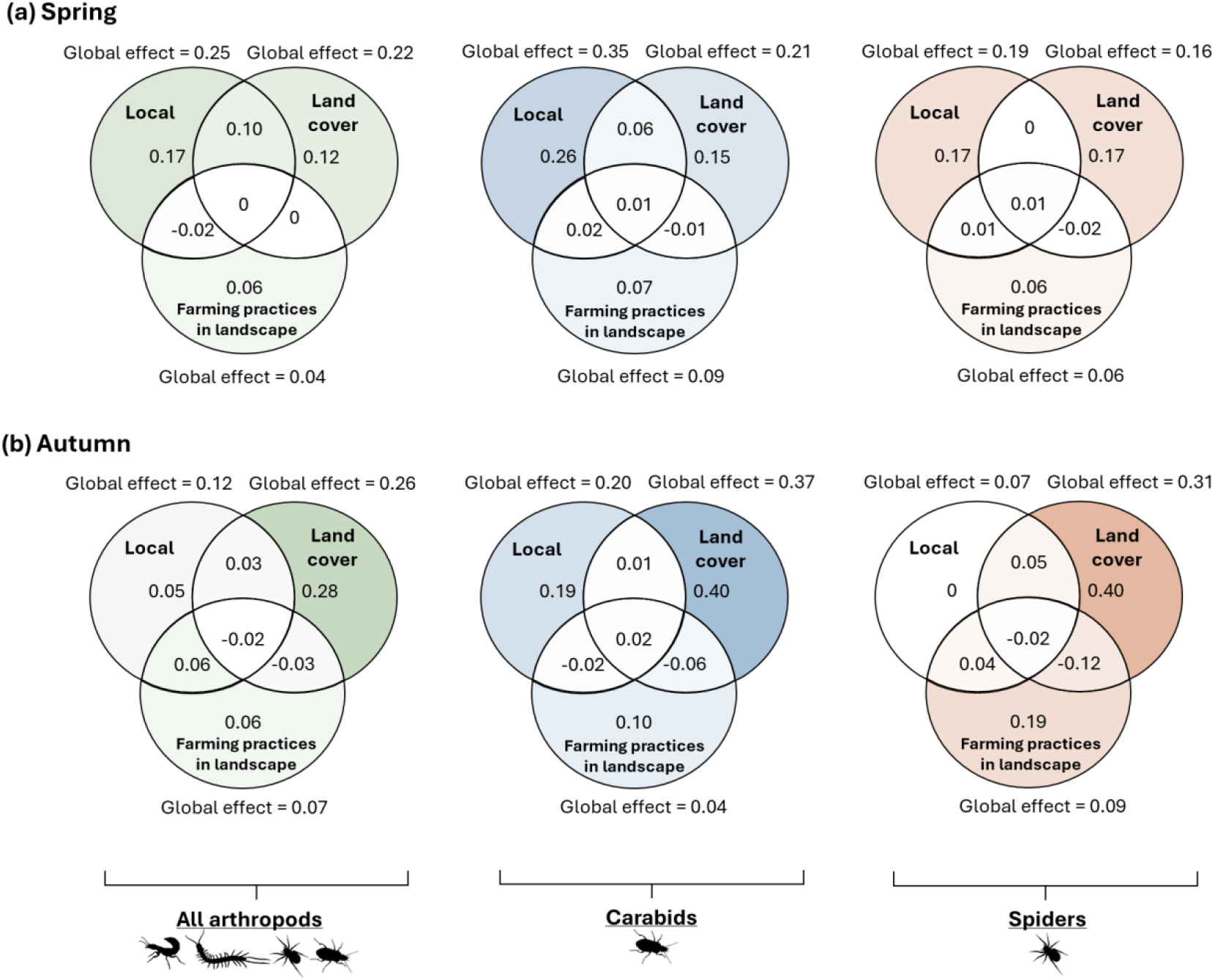
Venn diagram representing the mean proportion of variance of biological variables (a) in spring and (b) in autumn, explained by farming practices at landscape level parameters, local parameters, and land cover/resources parameters. The global effect corresponds to the sum of the values present in each of the circles and represent the proportion of the total variance explained by the group of parameters. The individual Venn diagrams for each biological variable are shown in Figures S5, S6.

In contrast to spring, landscape parameters, and especially farming practices, were most often selected in the models to explain biodiversity patterns (Figure 2b). Among the 18 models, local parameters were selected in 17 % of cases, land-cover parameters in 36 %, and farming practices at landscape level in 46 %. Parameters such as *LWI* (22.0 % of models), *NDVI-landscape* (55.6 %), and *proportion of grassland in the landscape* (22.0 %) showed recurrent effects on biological variables. The configuration of farming practices had also shown effects on certain biological variables through *d-low-I*. Farming practices at landscape level and farming practices combined with others environmental parameters explained a proportion of the variance for almost all the biological variables (Figures S5, S6), but only to a small extent (between 4 and 9 % in average) (Figure 3b). Most of the variance explained by farming practice at landscape level was explained alone and not in interaction with the other blocks. Local parameters explained between 7 and 20 % in average of the variance in biological variables, and land cover parameters between 26 and 37 %.

## 4. Discussion

Our study focused on the effect of both intensity and configuration of farming practices at landscape level on arthropod communities. By estimating the variance explained by farming practices at the landscape level compared to the variance explained by the other local and landscape parameters traditionally considered in studies, we are able to rank the anthropogenic effects.

We have shown that season directly changed parameters retained during model selection and their effects on arthropod communities. For the models associated with spring arthropod communities, parameters characterizing the state of the sampled grassland (i.e., *NDVI, temperature near the ground*, and *intensity of farming practices*) were selected more frequently than the landscape parameters. This can be explained by the fact that the arthropods found in grassland in spring are composed of organisms that could have taken refuge and overwintered there (Pywell et al., 2005; Sotherton, 1984). Their survival after winter may therefore depend above all on the environmental conditions provided by local grasslands themselves, as previously suggested (Sarthou et al., 2014). Conversely, for the models associated with autumn arthropod communities, the landscape parameters, and especially farming practices at the landscape level, were more often selected than local parameters. At this time of the year, some of the individuals that were in grassland for overwintering may have moved around the landscape in spring and summer, returning to the grassland after the harvest of crops. The identity and abundance of taxa present in grasslands in autumn depend on their ability to survive the environmental filters they undergo, linked for example, to the complexity of land cover and the intensity of farming practices (Lortie et al., 2004; Schweiger et al., 2005). This study demonstrates that the relationship between arthropods and agro-environmental parameters is modulated by the seasons. Consequently, conservation objectives and the means to achieve them need to be tailored to the phenology of these communities.

We showed effect of the intensity of farming practices at landscape level, and at several taxonomic levels. Given the difficulty of accessing this precise data over large landscape areas, the majority of studies dealing with their impact on biodiversity — and arthropods in particular — have until now been limited to discriminating between farming systems (i.e., organic vs. non-organic farming) (Brusse et al., 2024b; Petit et al., 2020), the data being freely accessible online. Whether on the level of arthropod communities or on the finer level of carabid beetle and spider communities, the intensity of farming practices at the landscape level is a driving force in the composition and diversity of permanent grassland arthropod communities, thus reinforcing the initial results observed by Brusse *et al*. (2024a). However, except a single effect on community composition, we did not find any effects of the configuration of practices on arthropods. This observation can be explained by two hypotheses: (i) arthropods are highly mobile, which makes them more sensitive to the general context of landscape intensity than to its specific configuration; (ii) given the high level of landscape connectivity in our study area, the amount of grassland and grass strips is already sufficient to maintain the species.

Our results revealed that farming practices at landscape level in our study area play a significant, albeit limited role, compared to local and land cover parameters. In our study area, local and land cover parameters therefore play a crucial role in arthropod preservation, confirming the observations of Puech *et al*. (2015) in another agricultural area but in a similar landscape context. The two areas are characterized by a gradient in the proportion of semi-natural habitats, generating a probable correlation between the land-cover gradient and the intensity gradient. It would be interesting to repeat these studies in areas with few semi-natural habitats to be sure of decorrelating the two gradients. Despite all this, our results do not invalidate the importance of integrating farming practices at landscape level in studies (Brusse et al., 2024b; Petit et al., 2020; Rusch et al., 2010). It remains complementary to that of the other environmental parameters, as shown by the low value of the interactions between the blocks of the variance partitioning. This result could also be an indication of the order of influence of environmental filters on community composition. According to a top-down logic, farming practice at landscape level would be the first and coarsest filter and would therefore have the least influence on community composition (Lortie et al., 2004).

We also highlighted a recurring effect of NDVI at the landscape level on arthropod communities, which was more marked than the effect of land-cover parameters classically recognized as drivers of arthropod communities (Estrada-Carmona et al., 2022; Marja et al., 2022; Tscharntke et al., 2021). The NDVI could therefore be more relevant for representing the quality of a landscape in the sense that it represents a physical dimension of the landscape (i.e. the state of the vegetation), whereas traditional parameters such as the quantity of grassland or crop diversity do not really reflect the physiological state of the vegetation in the landscape.

In terms of preserving grassland arthropod communities, our study suggests that multiscale management is necessary. Locally, dense and well-developed vegetation should be favored. The levers to be put in place could be a reduction in the intensity of mowing and grazing to preserve plant resources (Simons et al., 2014). On a landscape level, maintaining a spatiotemporal continuity of plant resources seems essential to provide all the resources necessary for arthropods throughout their life cycle (Estrada-Carmona et al., 2022). This continuity of resources can be ensured by diversifying the crops grown in the landscape. A reduction in the intensity of farming practices at the landscape level would also help to preserve arthropod communities. However, the ways in which farming practices can be managed at landscape level are not obvious and require further research. It seems that cooperation between farmers at landscape level is the best solution for implementing such landscape-level management of farming practices (Cong et al., 2014; Leventon et al., 2019; Marrec et al., 2022), but the dominant agricultural model does not encourage farmers to move in this direction. The successful implementation of coordinated management on a landscape level could depend on the implementation of non-individual incentives aimed at strengthening the coordination within groups of farmers (Villamayor-Tomas et al., 2019).

## 5. Conclusion

Our study revealed that spring and autumn grassland arthropod communities were influenced by different parameters of heterogeneity in agricultural landscapes. Whereas the spring communities are mainly influenced by local parameters, the autumn communities are mainly influenced by landscape parameters. NDVI (at both local and landscape scale) and the intensity of farming practices in the landscape, parameters that are rarely considered in studies, often influence arthropod communities. In addition, farming practices at landscape level explain a share of variance additional to that explained by the classic local quality and land-cover parameters. These results underline the importance of considering these parameters in studies, in addition to local and land cover parameters. In terms of research prospects, analyzing the interaction between the intensity of farming practices and other heterogeneity parameters would help us to understand how to manage agricultural landscapes to preserve biodiversity from intensive farming practices. The integration of other agricultural regions with different socio-economic contexts is also essential to verify the validity of our results on a larger scale.

## Acknowledgements

We would like to thank all farmers involved in this study for their interests in our survey, as well as the PNR of Lorraine. R.M and G.C. received funding from France AgriMer, under the framework of the Plan National de Recherche et d’Innovation 2021-2023 (project SEPIM).

## Statement of authorship

TB, RM, and GC conceived the study, RM and GC led the study, TB performed the study, JT collected biodiversity data, and all authors wrote the manuscript.

