## supplementary material for "Farming practices in the landscape have an impact on grassland arthropods, in addition to local conditions and land cover"

**Table S1**. List of all variables used to run the *microclima* model.

| Input data | Spatial resolution (m) | Temporal resolution | Source |
|---|---|---|---|
| Elevation | 25 | NA | IGN (https://geoservices.ign.fr/) |
| NDVI | 20 | monthly | Copernicus-Sentinel-2 (https://scihub.copernicus.eu/) |
| Forest height | 30 | NA | Global Ecosystem Dynamics Investigation (https://glad.umd.edu/dataset/gedi) |
| Crop height | 20 | monthly | Scientific literature and documents from French agricultural technical institutes |
| 2m temperature | 9000 | hourly | ERA-5 (https://cds.climate.copernicus.eu/cdsapp#!/dataset/reanalysis-era5-land?tab=overview) |
| 2m Relative humidity | 9000 | hourly | |
| 10m u/v-components of wind | 9000 | hourly | |
| Surface pressure | 9000 | hourly | |
| Cloud cover | 9000 | hourly | |
| Surface Solar Radiation Downwards | 9000 | hourly | |

**Table S2.** List of farming practice variables used to calculate the various agricultural practice intensity indices used in the article.

| Practice category | Variables | Description |
|---|---|---|
| *Pesticide treatment (the associated variables were duplicated for each type of pesticide treatment: herbicide, insecticide, fungicide)* | Number of treatments | Number of tractors passes over the plot to carry out a treatment, regardless of whether there are mixtures of active substances. |
| | Treatment frequency index (TFI) | Applied dose divided by the registered dose |
| | Total number of active substances | Total number of active substances applied to the plot. If the same active substances is applied several times, it is counted the number of times it has been applied. |
| | Active substances diversity | Number of different active substances applied to the plot |
| | Total number of active substances/Number of treatment | Ratio between the total number of active substances and the number of passages. This gives the average number of active substances per tractor pass over the plot. |
| | Mean time between each treatment | Average number of days between two pesticide treatments. We assume that the closer the treatments, the more intensive the variable |
| *Fertilization* | Number of fertilization | Number of tractors passes over the plot to carry out a fertilization |
| | Molecule diversity | Total number of molecules applied to the plot. If the same molecule is applied several times, it is counted the number of times it has been applied. |
| | Molecule diversity/Number of fertilization | Ratio between the molecule diversity and the number of passages. This gives the average number of molecules per tractor pass over the plot. |
| | Nitrogen unit | Total quantity of nitrogen applied to the plot (kg/ha) |
| | Phosphorus unit | Total quantity of phosphorus applied to the plot (kg/ha) |
| | Potassium unit | Total quantity of potassium applied to the plot (kg/ha) |
| | Proportion of N mineral fertilization | Proportion of total nitrogen added to the plot of mineral origin |

| | | |
|---|---|---|
| | Mean time between each fertilization | Average number of days between two fertilization. We assume that the closer the treatments, the more intensive the variable |
| *Soil tillage* | Total number of tillages | Number of tillage operations carried out on the plot |
| | Number of tillages with soil turning | Number of tillage operations carried out on the plot involving soil turning (e.g. ploughing) |
| | Diversity of tillage | Number of different tillage operations carried out on the plot. For example, ploughing, hoeing and decompacting the soil are different categories of tillage. |
| | Tillage depth | Depth of maximum tillage carried out in the plot |
| | Mean time between each tillage | Average number of days between two soil tillage. We assume that the closer the treatments, the more intensive the variable |
| *Mowing and grazing* | Number of mowing | Number of times vegetation has been cut in the plot. The value of 1 has been assigned to all crops because they are harvested only once. |
| | Mowing heigh | Height of vegetation left in the plot after cutting |
| | Livestock load | Number of animals weighted by their type in the plot. The value of 1 has been assigned to all crops because because they do not contain animals |

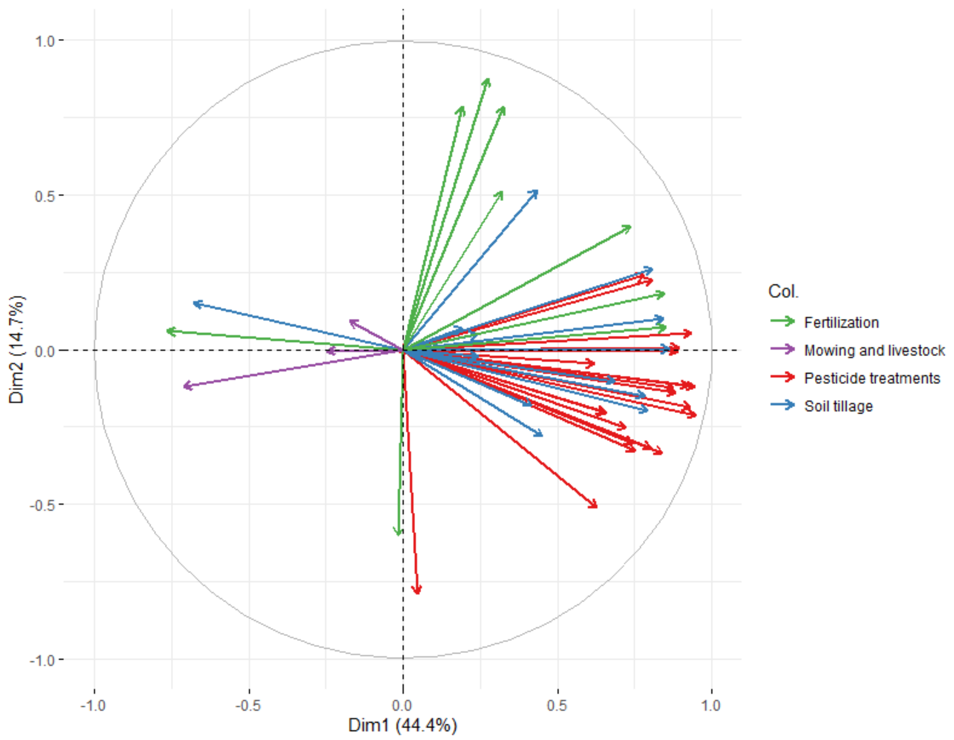


**Figure S1.** Principal Component Analysis (PCA) of each farming practice variable surveyed. Green arrows refer to fertilization variables, purple arrows to mowing and livestock related variables, red arrows to pesticide treatments variables, and blue arrows to soil tillage variables. Each variable consider in this PCA is detailed in Table S2.

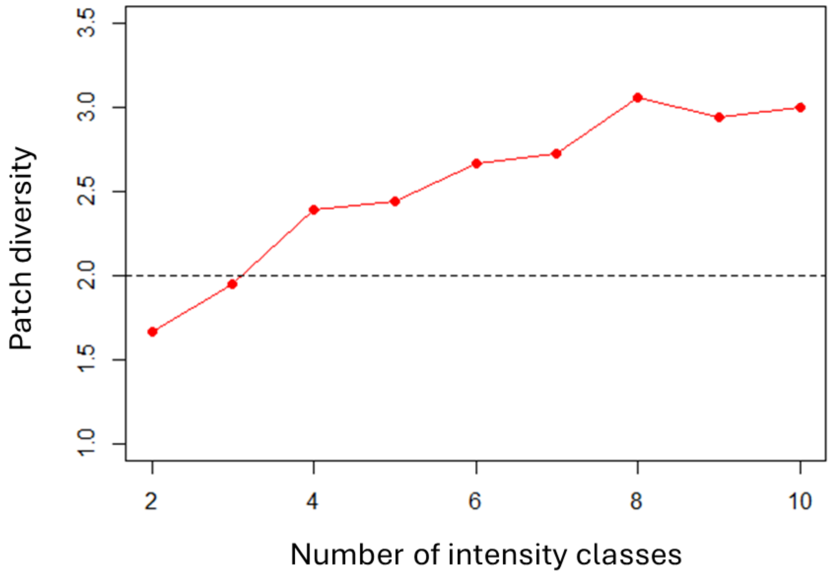


**Figure S2.** Evolution of patch diversity as a function of the number of intensity classes. Dividing the intensity into 8 intensity classes maximises patch diversity. The red line represents the patch diversity in the landscape linked to the intensity of agricultural practices. The dotted black line represents the diversity of patches in the landscape linked to land use (grassland vs. crops).

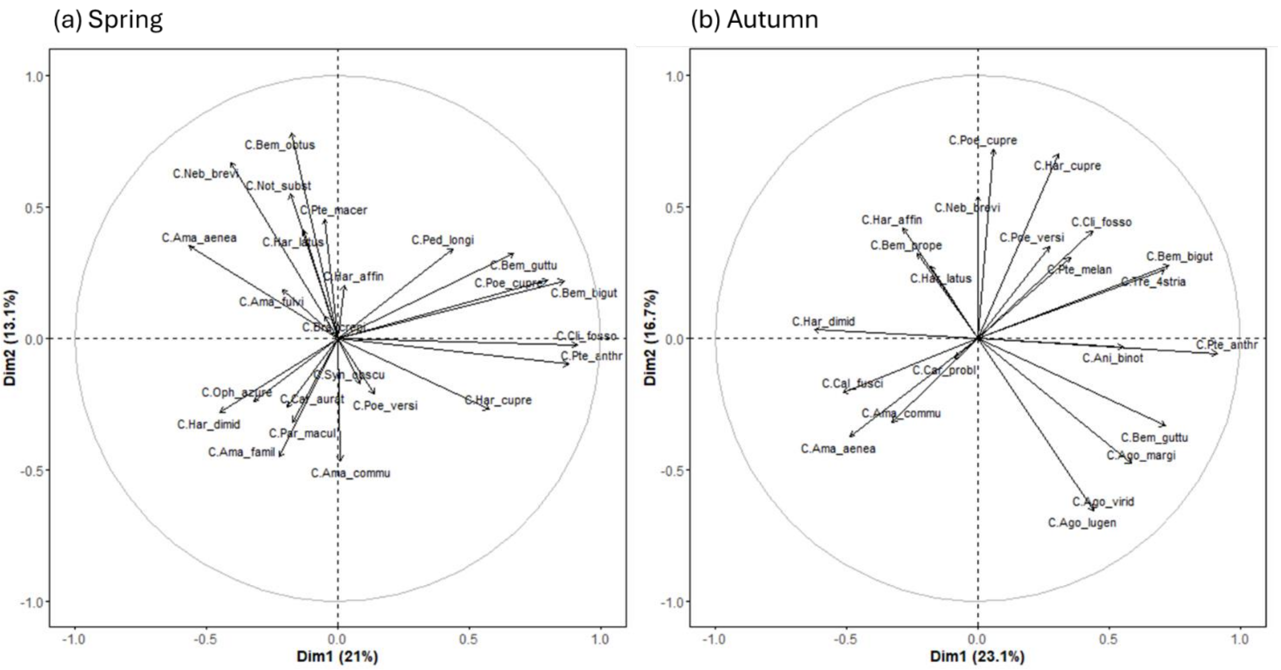


**Figure S3.** Results of principal component analysis (PCA) illustrating composition of carabid communities (a) in spring and (b) in autumn.

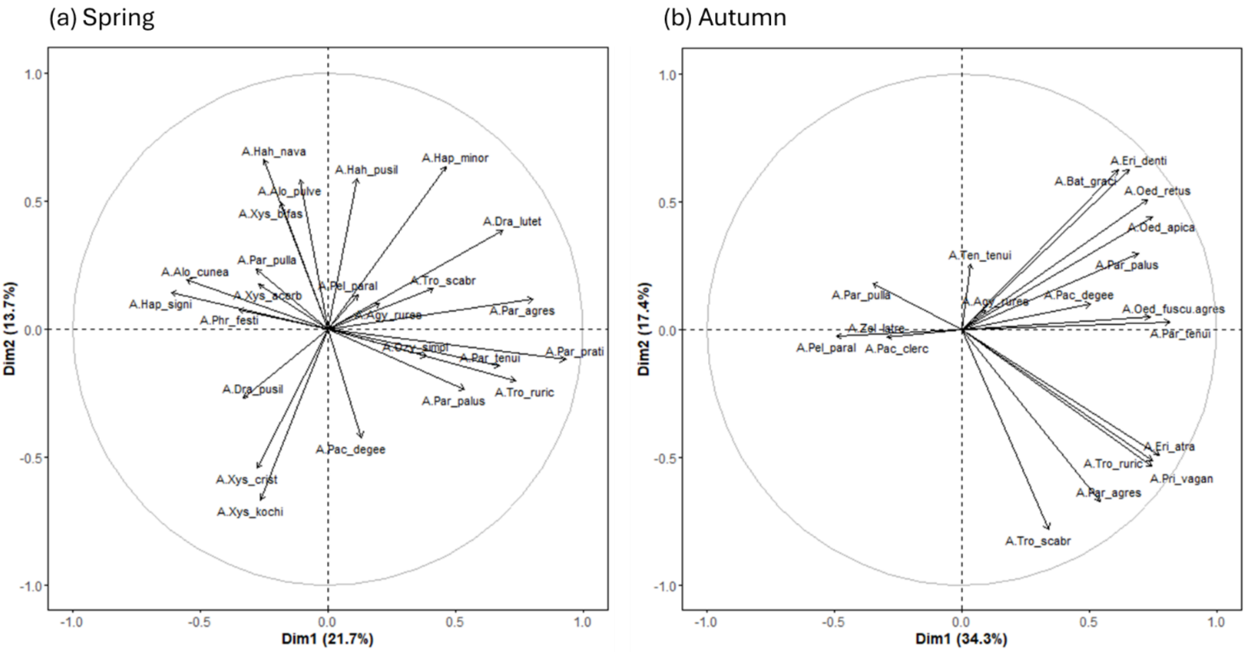


**Figure S4.** Results of principal component analysis (PCA) illustrating composition of spider communities (a) in spring and (b) in autumn.

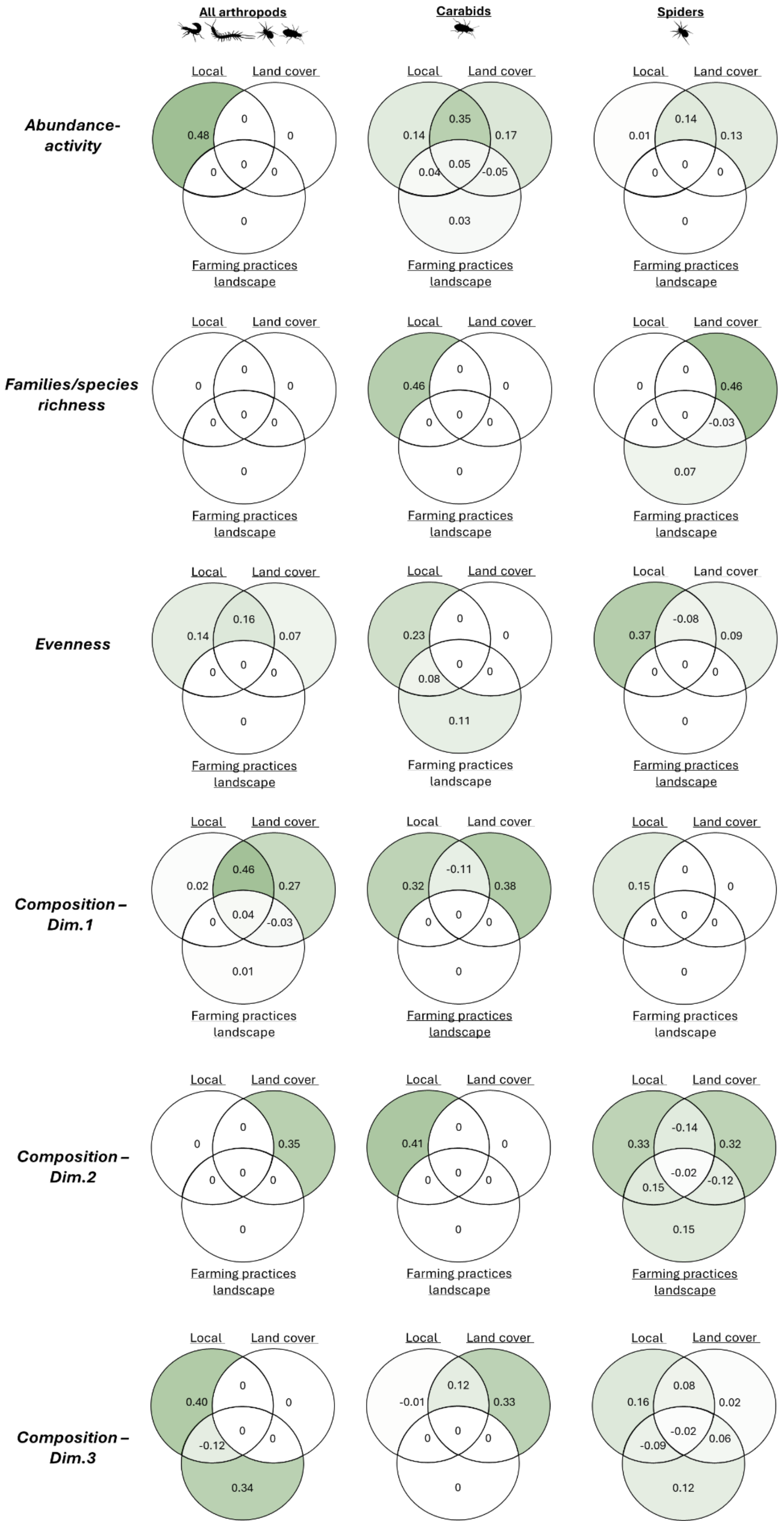


**Figure S5.** Individuals Venn diagrams representing the proportion of variance of each biological variables in spring, explained by farming practices at landscape level parameters, local parameters, and land cover/resources parameters.

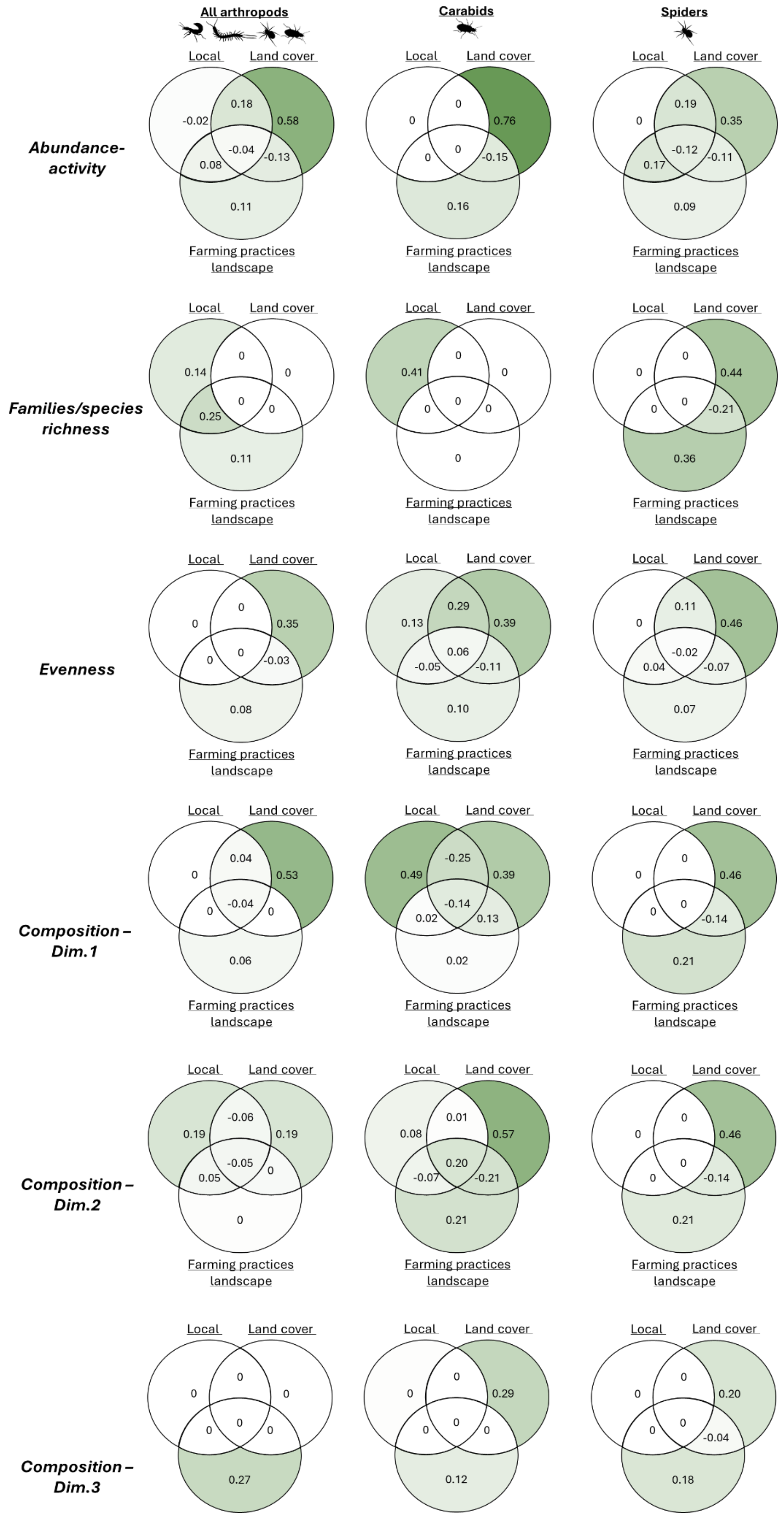


**Figure S6.** Individuals Venn diagrams representing the proportion of variance of each biological variables in autumn, explained by farming practices at landscape level parameters, local parameters, and land cover/resources parameters.
